# Lichen Growth Form Effects on Rock Weathering in Cold Biomes

**DOI:** 10.64898/2026.06.22.733894

**Authors:** E. N. Ciric, I.K. de Jonge, R. Liu, J. H. C. Cornelissen, P. Convey, S. Bokhorst

**Affiliations:** Amsterdam Institute for Life and Environment (A-LIFE), Vrije Universiteit Amsterdam, 1108, 1081 HZ Amsterdam, The Netherlands; British Antarctic Survey, Natural Environment Research Council, High Cross, Madingley Road, Cambridge CB3 0ET, United Kingdom; Department of Zoology, University of Johannesburg, PO Box 524, Auckland Park 2006, South Africa; Millennium Institute – Biodiversity of Antarctic and Sub-Antarctic Ecosystems (BASE), Santiago, Chile; Cape Horn International Center (CHIC), Puerto Williams, Chile; School of Biosciences, University of Birmingham, Edgbaston, Birmingham B15 2TT, United Kingdom

**Keywords:** Antarctic biodiversity, plant ecology, cryptogam traits, terrestrial ecology, effect traits, interspecific variation

## Abstract

Rock surface weathering is a critical element in the process of early soil formation, in which lichens are thought to play a significant role. Crustose lichens, with a large area of rock-surface contact, are generally considered more influential in rock weathering, while foliose and fruticose growth forms, with more developed three-dimensional structure and less rock-surface contact, are rarely considered in this context. Here, we test the extent to which all three growth forms contribute to granitic rock surface weathering in Maritime Antarctic ecosystems, by quantifying rock hardness beneath foliose (n = 2 species), fruticose (n = 2) and crustose lichens (n= 5). Our data confirm that foliose lichens reduced rock surface hardness by 9%, to a lesser extent than crustose and foliose lichens (40% and 31% reduction, respectively). To disentangle whether these effects result from lichen-induced weathering or lichen preference for pre-weathered rock, we also analyzed a dated deglaciation sequence on granitic rocks from the Morteratsch Glacier forefield in the Swiss Alps. At this location, the impact of crustose lichens on rock substrate hardness generally increased with time since exposure from glacial retreat and with lichen thallus size. We conclude that lichen presence on rock surfaces significantly reduces rock hardness, with crustose lichens having a greater impact than foliose and fruticose forms, highlighting the potential role of lichens of all three growth forms in driving substrate breakdown and shaping early-stage ecosystem processes in polar and alpine regions.

## Introduction

Lichens are an important component of terrestrial biodiversity in cold biomes (Matveyeva and Chernov 2000; Nimis et al. 2018), where they perform important ecosystem functions including rock weathering (Adamo and Violante 2000; Cornelissen et al. 2007). Lichen growth form is an important trait influencing their contributions to ecosystem processes and biodiversity (Asplund and Wardle 2017). As early colonizers of exposed terrestrial habitats, their presence and growth form may influence rock weathering (Lindsay, 1978; Jie and Blume 2002). Rock weathering is a critical contributor to the early stages of soil formation, however, lichen species contribute differentially to this process (Chen et al 2000). Lichen rock weathering is primarily driven by the release of acidic ‘lichen compounds’ and hyphal growth into micro-fissures in the rock surface (Jie and Blume 2002). Among the various growth forms that characterize lichens, crustose lichens — which characteristically have about 50 % direct tissue contact with the rock surface — appear the most likely to contribute significantly to rock weathering and have received the most research attention (Wilson and Jones 1983). However, foliose and fruticose lichens can also be firmly attached to rock substrata and have extensive distributions in many cold biomes (Adamo and Violante 2000; Øvstedal and Smith 2001) but, compared with crustose species, few studies are available on the rock weathering effects of these lichens (Chen et al. 2000). The western Antarctic Peninsula region (Maritime Antarctic) hosts around 350 lichen species representing all three growth forms (Øvstedal and Smith 2001). Although rock weathering by crustose lichens has been demonstrated in this region (McCarroll and Viles 1995; Matthews and Owen 2008; Guglielmin et al. 2012), research has yet to address any contribution of foliose and fruticose lichens despite their regional abundance (Holdgate 1977; Peat et al. 2007).

Lichens can enhance rock weathering chemically via their natural secretion of acidic lichen compounds (Chen et al. 2000), or physically by the action of thallus contraction and expansion or through hyphae infiltrating and enlarging micro-fissures in the rock surface (Syers and Iskandar 1973; Purvis et al. 2013). Crustose lichens form an intimate association with the underlying rock substratum and there is visible evidence of hyphae growing into rock surfaces (Adamo and Violante 2000). These hyphae are often well enmeshed into the substratum, making physical removal difficult and damaging to both the lichen and the rock (Armstrong and Bradwell 2010). Crustose lichens can colonize and weather a wide variety of substrata including different rock types, concrete, leather materials, old timber, bones and living organisms such as trees and mosses (Kappen 2000).

Contrasting with crustose species, foliose lichens have a more leaf-like shape, while fruticose lichens are shrub-like, often with multi-branched thalli. Both of these growth forms attach to the substratum at a single anchoring point (Asplund and Wardle 2017). This lichen anchoring point provides direct influence on the rock through hyphal attachment (Ascaso and Wierzchos 1994) and through leaching of acids and organic compounds (Syers and Iskandar 1973). Although the direct effects of foliose and fruticose lichens on rock weathering are widely assumed to be smaller than those of crustose lichens, they are unlikely to be zero and therefore could still contribute to habitat formation in the nutrient-and mineral-deficient ecosystems typical of polar biomes. However, no empirical data are available to quantify their weathering effects.

Quantifying rock weathering by lichens is challenging given the slow growth rates of these organisms, the typically hard nature of rock substrata and the generally slow rates of geological and ecological processes in polar biomes. Lichen impacts on rock surfaces most likely commence upon first establishment but, with varying radial growth rates of 0.01 - 0.9 mm year^-1^ measured in Antarctica (Sancho et al. 2007), it may take several decades before quantifiable surface areas are weathered. A rapid and practicable approach for assessing lichen influence is to quantify rock hardness at the site of lichen growth and compare this against adjacent lichen-free rock surfaces (Matthews and Owen 2008). Rock surfaces in Antarctica typically show lower rock hardness values over time since exposure, i.e., the surface is weakened by environmental weathering (Guglielmin et al. 2012, Kanamaru et al. 2018). However, whether any differences in rock hardness thereby detected result from active lichen weathering or a preference for lichen establishment on already weathered rock surfaces requires clarification. To address this cause-and-effect question, complementary approaches are required, such as quantifying lichen rock weathering effects across dated surfaces. If lichens actively weather the rock surface, their effect on rock hardness should be greater on older surfaces with a longer period of lichen presence. Furthermore, lichen size is a reasonable indicator of age, especially in radially growing crustose lichens (Armstrong 2011) and, therefore, larger lichens should have a stronger impact on rock hardness than smaller ones, which will generally have been present for a shorter period of time.

In this study, we compared the influence of different lichen growth forms on rock hardness, as a relative measure for weathering in the Maritime Antarctic (as applied by Guglielmin et al. (2012)) beneath nine common lichen species. The lichen species selected included crustose (n = 5 species), foliose (2 species) and fruticose (2 species) growth forms. Further, to address the cause-and-effect question related to lichen presence and rock hardness, we quantified lichen effects on relative rock hardness on similar granitic substrates across a dated glacial chronosequence (123 years) in the Swiss Alps, including measurements of four crustose species. We hypothesized that: (1) crustose lichens will have a greater impact on rock hardness than foliose and fruticose lichens, (2) lichen impacts on rock hardness are a result of active weathering processes and, therefore, will be greater on older surface exposure rocks than younger ones, and (3) in line with the time since deglaciation, lichen size (as a proxy for age) will be positively related to reductions in rock hardness.

## Methods

### Measurements

Two study regions were selected based on their similarity in rock type (granitic) and hosting terrestrial ecosystems with high cryptogam biodiversity. Antarctic lichen species’ effects on relative rock hardness were quantified on East Lagoon Island, which is located in Ryder Bay (Adelaide Island) west of the Antarctic Peninsula mainland (67°35′34″S 68°14′06″W) and close to the British Antarctic Survey’s Rothera Research Station. The local climate is typical of the maritime Antarctic (Bokhorst et al. 2007; Convey et al. 2018). The island’s terrestrial habitats are characterised by loose boulder beaches and an interior consisting of granitic outcrops up to 50 meters above sea level (m a.s.l.), as well as shallow rocky valleys (Guglielmin et al. 2012). East Lagoon Island has scattered persistent snow patches in summer and meltwater channels with mosses and some grass, as well as fellfields colonised by various lichens (Convey and Smith 1997). Twenty-seven lichen species are currently recorded from East Lagoon Island, including crustose, foliose and fruticose growth forms (ASPA Management Plan 2021). We selected nine abundant lichen species to quantify their impact on rock hardness, including crustose lichens (*Buellia frigida, B. russa, Lecidea atrobrunnea*, *Rhizocarpon geographicum, Xanthoria elegans*), fruticose lichens (*Pseudephebe minuscula* and green non-melanised and black melanised variants of *Usnea antarctica*) and foliose lichens (*Umbilicaria antarctica*, *U. decussata*) (Øvstedal and Smith 2001). *Usnea antarctica* (green variant)*, R. geographicum* and *X. elegans* were sampled in well-shaded valleys on East Lagoon Island, while all other species were sampled in sun-exposed areas.

Rock hardness was measured applying the Schmidt Hammer method, using a spring-loaded hammer (Proceq SilverSchmidt PC Type N) that quantifies rock hardness to indicate its surface weathering through its rebound value (Matthews and Owen 2008; Guglielmin et al. 2012; Matthews and Winkler 2022). Rock hardness values were converted from R-values using an exponential calibration function (Viles et al. 2011). However, this conversion produced an almost linear relationship within the observed range of R-values (R² = 0.998). Consequently, analyses conducted using raw R-values or converted hardness values gave equivalent results. Rock hardness beneath lichens was obtained through two consecutive hammer strikes on the same spot for 25 individual lichen specimens per species, with the value of the second strike recorded. The repeat strike was made to reduce interference from any remaining lichen debris at the strike location (Guglielmin et al. 2012; Matthews and Winkler 2022). The sites were selected based on the abundance of the lichen species most commonly observed at each. To obtain hardness of the rock surface without lichen presence, we selected a nearby point (<5 cm distance) with the same rock type, free of visible organic matter. This sampling regime resulted in a split plot design with matched bare rock and lichen-covered sampling sites (n = 25 per species). Fruticose and foliose lichens were removed by twisting the lichen from its attachment point before hardness measurements to avoid hammering through dense lichen matter. Crustose lichens were left in place as removal would have only been possible by scraping, thereby affecting the rock surface. The presence of the crustose lichen could possibly add a flexible layer over the rock and affect the rebound value. Because two blows were always performed and due to the lichen’s flat thallus, the presence of a crustose lichen is assumed not to have influenced the rebound value, a method employed in other studies (Matthews and Owen 2008). Even with ten to thirty repeated strikes on the same crustose lichen, rebound values plateau and are not the same values as those of bare rock (Guglielmin et al. 2012).

To avoid any impact of surface moisture on rock hardness, measurements were taken on dry days with visibly dry rock surfaces (Matthews and Owen 2008). Additionally, to avoid lower values caused by structural variations, surfaces that were chipped, cracked, unstable, or otherwise atypical in appearance were avoided. Sampling was only carried out on horizontal or near-horizontal surfaces, and the boulders were not cleaned or scraped prior to sampling as such pre-treatment (although suggested by the manufacturer) was not applied in this case because polishing would impact or completely remove the weathered layer (Viles et al. 2011). All measurements beneath lichens were taken as close to the center of the thallus or attachment point as possible. Rock hardness values (R-values) lower than 40 were removed from the dataset, as this number is below the typical R-value of 50-60 for granitic rocks (Goudie 2006) and are considered to reflect displacement of the hammer tip or the rock surface crumbling or fracturing upon impact.

To test if rock hardness, a proxy for weathering (Matthews and Owen 2008, Viles et al. 2011), is influenced by lichen presence rather than reflecting a preference by lichens to establish on weathered rock, we compared lichen influence on rock hardness along a known dated age (weathering) sequence of a retreating glacier. If lichen presence is influential, active lichen weathering processes should result in lower rebound values over time. Several studies in alpine and polar environments including glacier moraines and shorelines have used the Schmidt Hammer to quantify the relationship between rock weathering and the length of time the surface has been exposed to weathering (Matthews and Shakesby 1984, Matthews and Owen 2008, Viles et al. 2011). It is important to note that evaluating a surface’s year of exposure based on Schmidt hammer rebound values will always be a relative measure based on reference rebound values (Rune and Sjåstad 2000).

The Swiss Alps were chosen for this part of our study due to the high quality and availability of archival maps and aerial photography from Switzerland’s national mapping agency (Swisstopo). We selected the forefield of Morteratsch Glacier near Pontresina, Switzerland, in the Bernina Alps (46°25′47″N 9°55′58″E) due to the granitic composition of the rocks in its forefield, its ease of access, and relatively low altitude (1950-2100 m a.s.l.) while still qualifying as a cold area. This is a northerly-flowing alpine glacier, surrounded by granite substrata of a similar type to that found on Lagoon Island. The valley has a mean annual air temperature of -9°C, is protected from wind, and winter snow cover occurs between October and April (Climate-Data.org 2021). Like other glaciers in the Swiss Alps, the Morteratsch Glacier has receded significantly since the end of the Little Ice Age (Egli et al. 2006) and this retreat has been documented since the 1850s (Burga et al. 2010). We defined eight sampling sites with known deglaciation occurring at 2022, 1998, 1971, 1955, 1930, 1917, 1896 and 1875. At each site, we selected 10 individual boulders (n = 10 replicate sampling sites at each deglaciation stage) over 1 m in diameter with a typical appearance at each site and quantified relative rock hardness in the same manner as described above. This part of the study included four crustose lichen species, *Aspicilia cinerea, Aspicilia* sp., *Rhizocarpon geographicum* and *Sporastatia testudinea,* of which *R. geographicum* (sampled in both regions) also served as a check for comparability between the two sampling regions. We applied 50 hammer strikes per boulder (n = 5-20 per species and neighbouring control values). For each lichen, we measured thallus diameter (mm) as a proxy for lichen age, in order to quantify if this influenced any within-species lichen effect on rock hardness.

### Statistical analyses

For the Antarctic data, two-way analysis of variance (ANOVA) was used to test for effects of lichen presence on relative rock hardness for growth form and lichen species separately. To address the existing variation in lichen effects on relative rock hardness within and between species across sampling sites, we calculated a log response ratio (LRR) based on the individual hammer strikes of each paired lichen and bare rock (y = log (x_1_/x_2_)), where *x*_2_ is the bare rock (control) strike and *x*_1_ is the lichen (presence) strike. This approach allows for a standardized comparison of the relative effect of lichens on rock hardness, independent of the hardness of the control rock. Species mean LRRs were calculated from 25 replicate pairs. A one-sample T-test was carried out on the LRR data to identify whether species had a significant influence on relative rock hardness.

For the Swiss data, analysis of covariance (ANCOVA) was used to quantify the effect of lichen presence, species identity and time since deglaciation on the LRR of rock hardness values, with thallus size as covariate. Size was categorized into two groups—> 10 mm and < 10 mm thallus diameter—based on an approximately equal distribution of data points between the two classes. Before conducting the ANCOVA, we tested the assumption of homogeneity of regression slopes with a linear model. To further investigate the relationships between LRR and covariates (year and size), we used a linear model to evaluate potential differences in slopes. Pairwise comparisons with adjusted P-values (to account for multiple comparisons) were then made to quantify whether significant differences existed between species and year after accounting for the covariate size. A linear model was used to identify if lichen presence, year of exposure and the combination of these two factors had statistically significant effects on reduction in rock hardness. All analyses were carried out in R (R Core Team 2024).

## Results

### Impact of lichen growth form on relative rock hardness

The converted hardness values of bare rock at the East Lagoon Island study site had a mean value of 148. Converted hardness was, on average, significantly lower beneath lichens, with a mean of 103 (Table 1, Fig. 2a). However, significant differences were observed only for three crustose species, *B. russa*, *B. frigida* and *X. elegans*. The log response ratio of rock hardness between presence and absence of lichens indicated that 8 of the 10 examined lichen species reduced rock hardness (Fig. 2b). Crustose lichens (mean 40 %) and foliose lichens (mean 31 %) reduced rock hardness the most, while fruticose lichens (mean 9 %) had smaller impacts (Fig. 2, Table 1; Supplementary Table S1).

**Fig. 1a.**
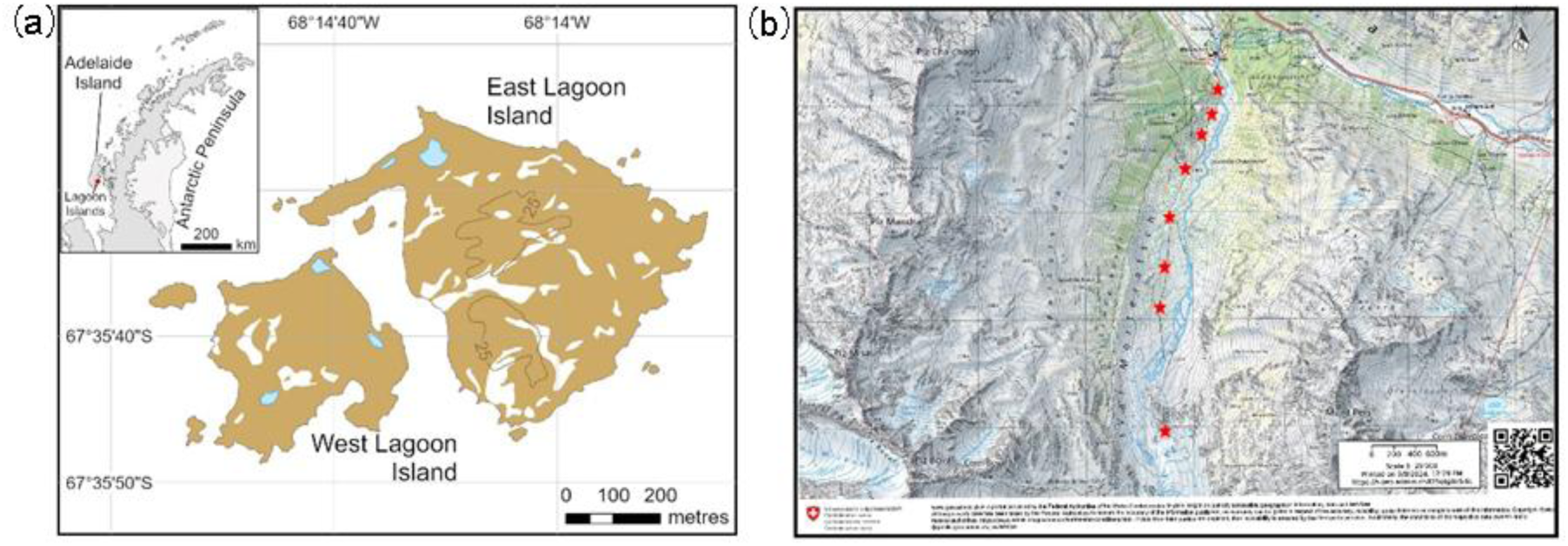
The Lagoon Islands in Ryder Bay, off the west coast of the Antarctic Peninsula. All samples were taken on East Lagoon Island. Map produced by the BAS Mapping and Geographic Information Centre, with data from the SCAR Antarctic Digital Database, 2024. Figure 1b. Sampling sites along the forefield of Morteratsch Glacier in the Bernina Alps, Switzerland, based on deglaciation age. Each red star corresponds to the estimated year that the area became deglaciated, from south to north: 2022, 1998, 1971, 1955, 1930, 1917, 1896, and 1875, the latter being the furthest from the current glacier terminus.

**Fig. 2.**
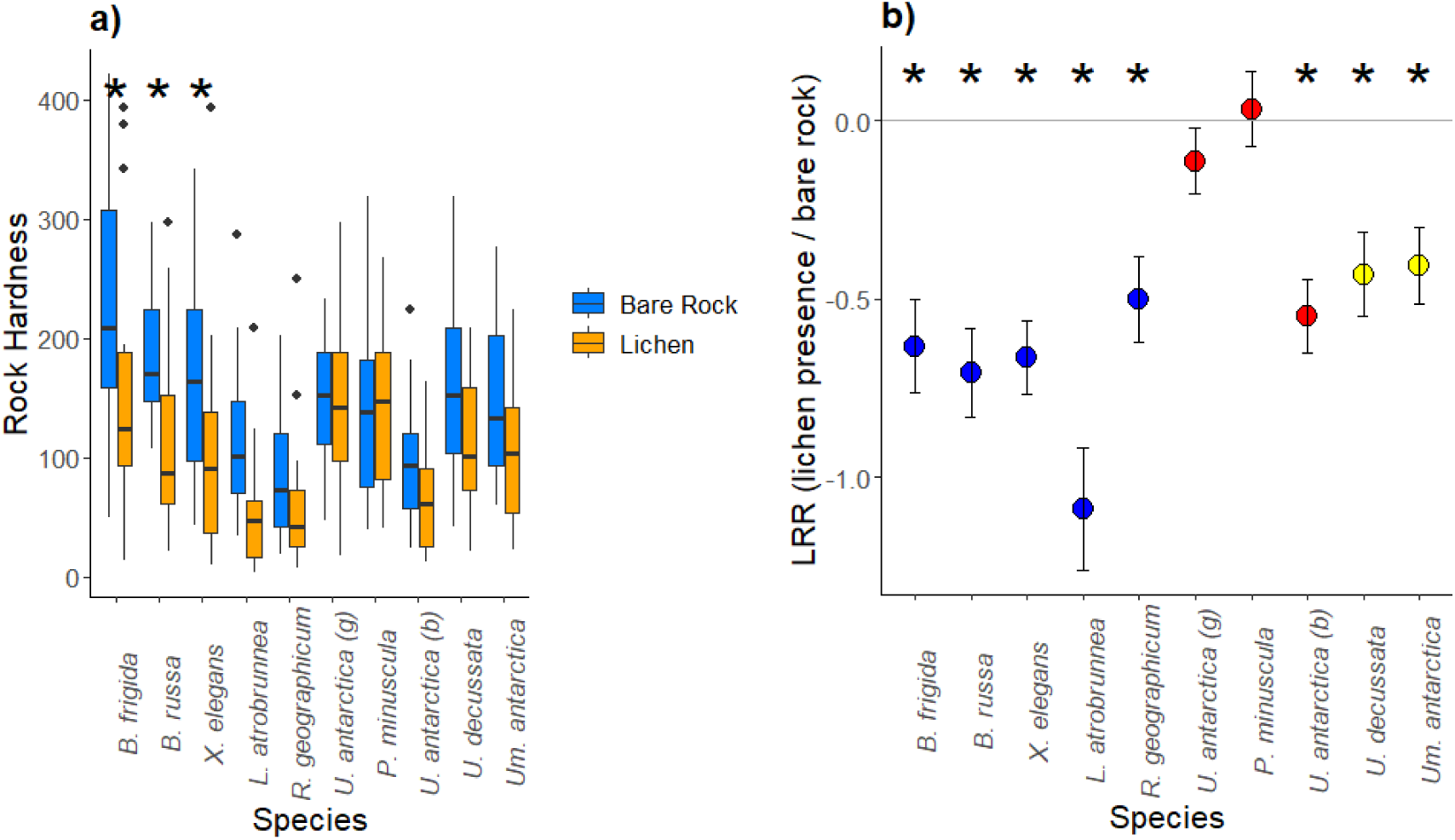
The impact of different lichen species on rock hardness on East Lagoon Island, Maritime Antarctic. a). Box-plots of mean hardness of rock surface (Bare Rock) and beneath Antarctic lichen species (Lichen). b) Log response ratio of hardness values between lichen presence and bare rock. Values below zero indicate a lower rock hardness beneath lichens compared to bare rock. * indicates significant difference (Tukey HSD P < 0.05) in relative hardness between bare rock and lichen presence. Mean values are based on 25 replicate measurements per species, with lines indicating standard error. Species are grouped by growth form and ordered from high to low rock hardness. Crustose: *Buellia frigida, B. russa, Xanthoria elegans, Lecidea atrobrunnea, Rhizocarpon geographicum*; fruticose: *Usnea antarctica* (green and black variants, referred to as (g) and (b), respectively), *Pseudephebe minuscula*; foliose: *Umbilicaria decussata* and *U. antarctica*.

**Table 1.** ANOVA output (F- and P-values) of the impact of Antarctic lichen species and growth form on rock hardness. Sample sizes were n = 10 for species and n = 3 for growth form.

|  | Species |  |  | Growth form |  |  |
| --- | --- | --- | --- | --- | --- | --- |
|  | Df | F | P | Df | F | P |
| Lichen Presence | 1 | 54.4 | < 0.001 | 1 | 43.6 | < 0.001 |
| Species/Growth form | 9 | 14.6 | < 0.001 | 2 | 0.52 | 0.594 |
| Lichen Presence × SP/GF | 9 | 2.31 | 0.015 | 2 | 5.48 | 0.004 |

### Chronosequence of rock hardness

The lichen impact on rock hardness increased significantly with age of exposure and lichen thallus size (Table 2, Figs. 3-5). Hardness values were more variable beneath lichens (20 – 452) compared to bare rock (101 - 335). Relative hardness of bare rock declined with time since exposure (R^2^ = 0.16), but this decline was stronger over time for rocks with lichens (R^2^ = 0.21). Changes in rock hardness over time since deglaciation differed significantly between study species (Fig. 4). Rock hardness declined by 1.04 y^-1^ in *Sporastatia testudinea,* 0.417 y^-1^ in *A. cinerea,* and 0.407 y^-1^ in *Aspicilia* sp. Only *R. geographicum* had a negative slope of -0.166 y^-1^, indicating that rock hardness did not decline with this species. ANOVA confirmed that the output was statistically significant for both year and species, but not their interaction, so indicating that year and species effects were consistent (Table 3). Relative rock hardness also declined with lichen thallus size for these three species, although with large variation (Fig. 5).

**Fig. 3.**
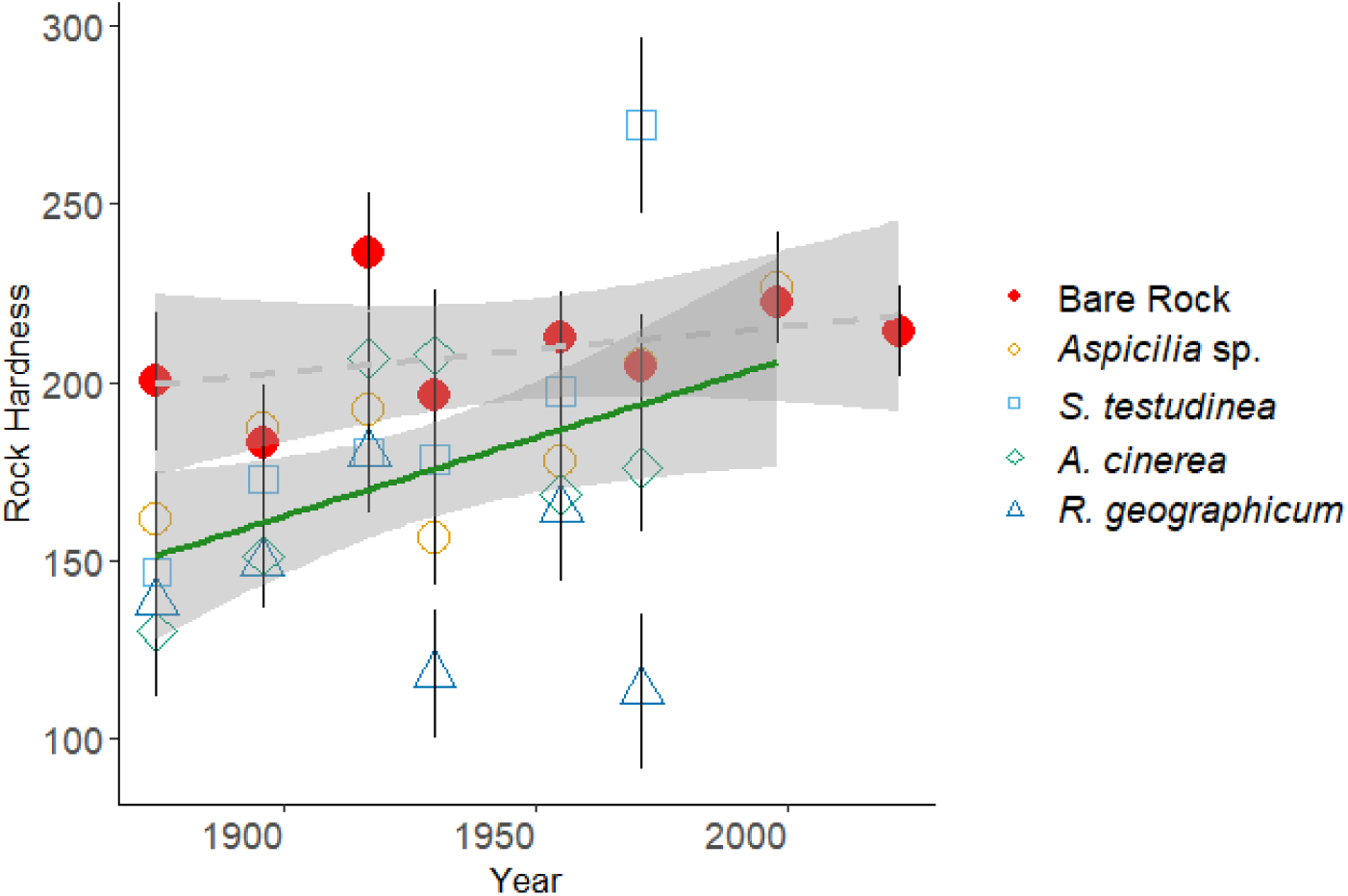
Rock hardness and lichen influence along a dated chronosequence of glacial retreat at the Mortserach Glacier in the Swiss Alps. Note that multiple lichen species, shown by different symbol type and colour, were present at most sampling sites, excepting the 2022 site, where no lichens were visible on the rock surface. Trend lines indicate change in rock hardness over time for bare rock (dotted line) and beneath lichens (solid line). Symbols are the mean of 10 replicate samples at each chronosequence point (with SE as error bars), and 95% confidence intervals are represented by grey shading. ANOVA output is presented in Table 3.

**Fig. 4.**
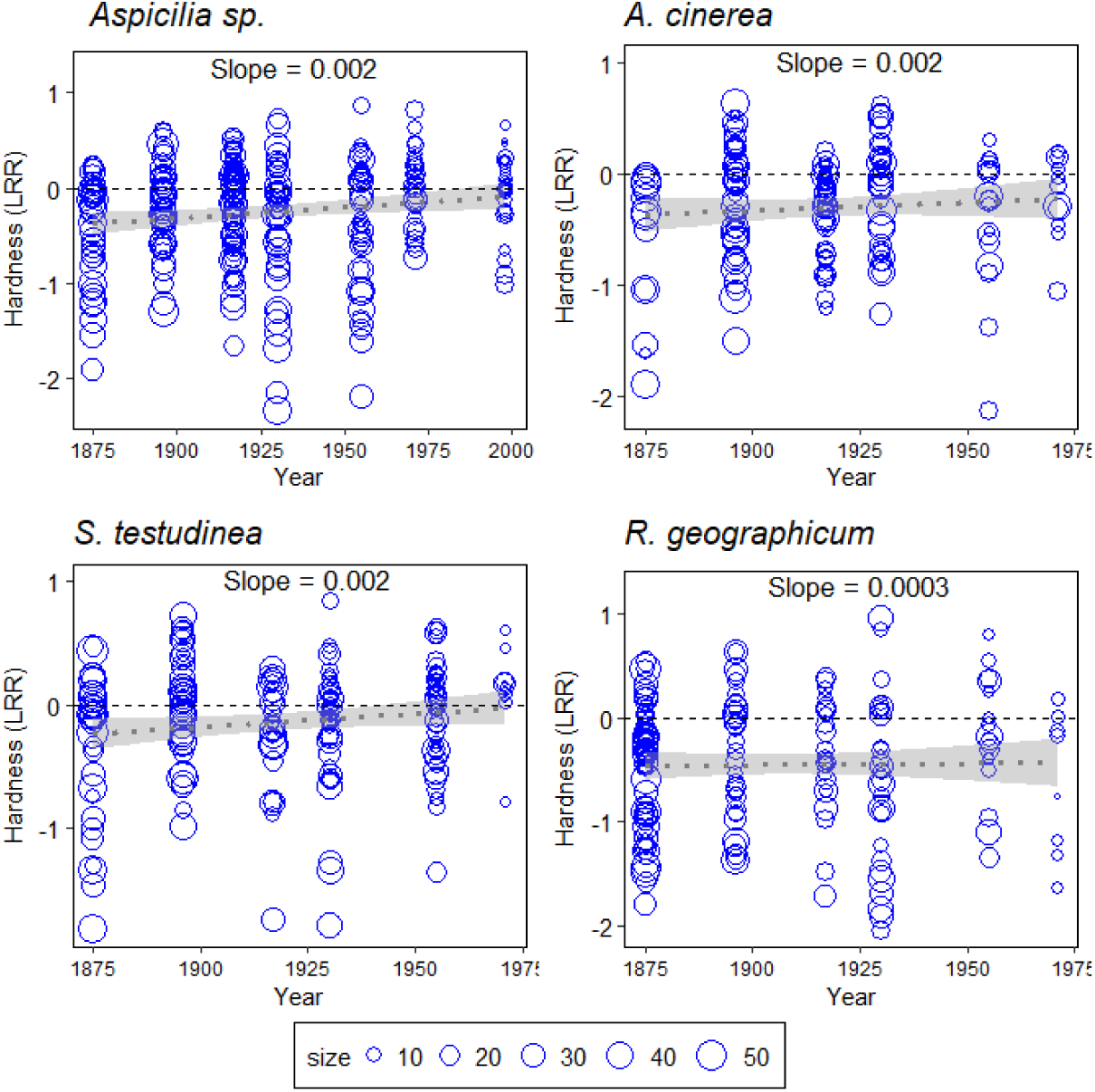
Rock hardness beneath four lichen species quantified along a dated glacial chronosequence in the Swiss Alps. Negative log response ratios indicate lower rock hardness beneath lichen compared to bare rock. The dotted trend line indicates the overall pattern over time since deglaciation. Ninety-five percent confidence intervals are represented by grey shading. Each symbol represents an individual lichen thallus (90-150 per panel) with symbol size reflecting thallus diameter.

**Fig. 5.**
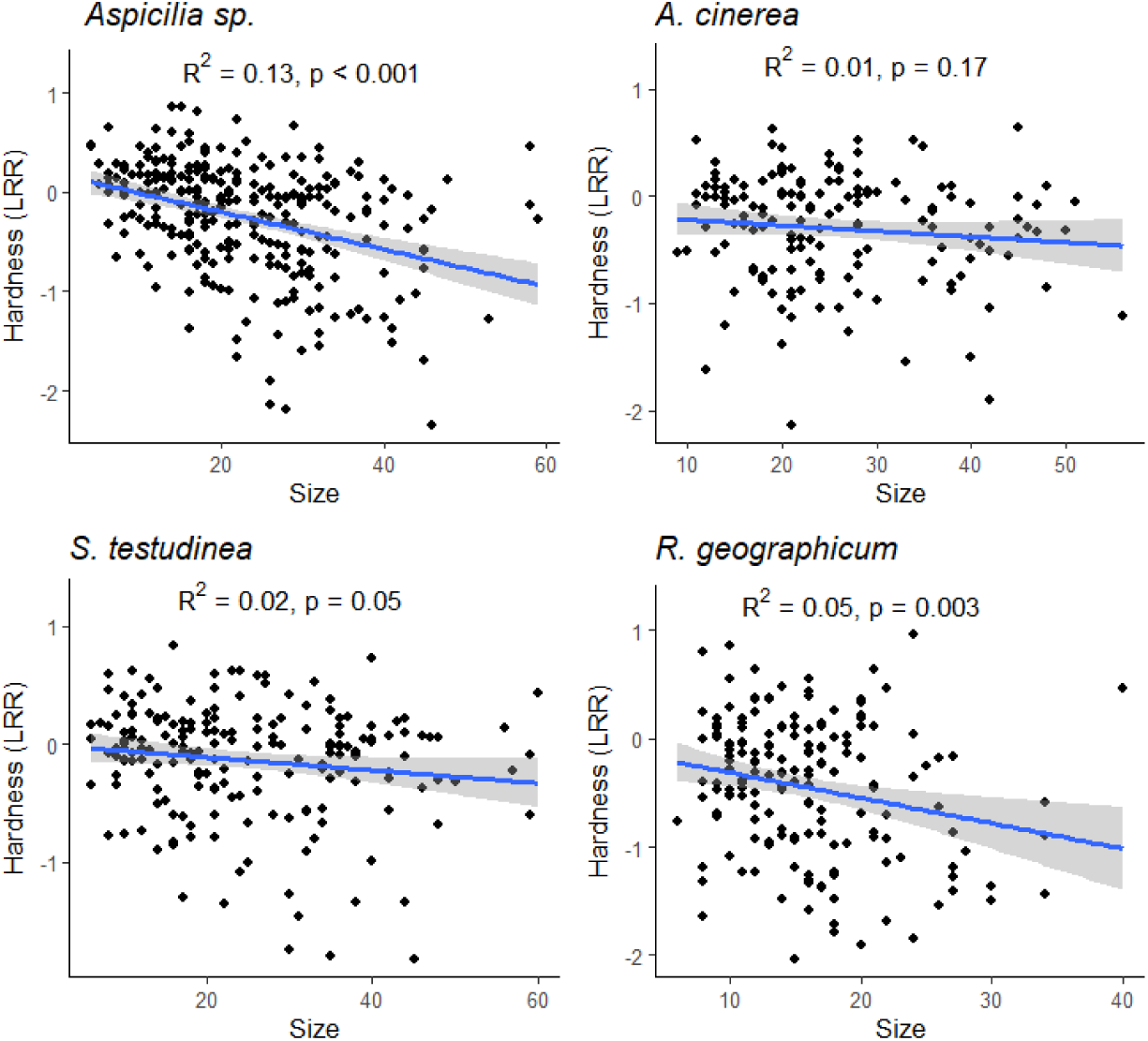
Lichen thallus size influence on rock hardness. The log response ratio of rock hardness with lichens present compared to absent is shown for different sized thalli separately for the four studied lichen species. Each data point is the log response ratio measured across a dated chronosequence in the Morteratsch Glacier forefield of the Swiss Alps. There are 90 to 150 values per species. Values of LRR < 0.0 indicate lower rock hardness. Ninety-five percent confidence intervals are represented by grey shading.

**Table 2.** Output from an ANCOVA of the log response ratio (LRR) of rock hardness in the presence or absence of lichens. The model tests the effect of lichen species, time since exposure and lichen size class on LRR. There are two lichen diameter classes, > 10 mm and < 10 mm. Data were obtained from a dated chronosequence in the Swiss Alps, with LRR as dependent variable and species (n = 4), year (n = 7) and thallus size as independent variables.

| Variable | Df | F value | P Value |
| --- | --- | --- | --- |
| Intercept | 1 | 3.4 | 0.07 |
| Species | 3 | 13.8 | < 0.001 |
| Year of Exposure | 6 | 3.4 | 0.003 |
| Size Class | 1 | 49.3 | < 0.001 |

**Table 3.** Output of ANOVA of a linear model testing the impact of all four species (*A. cinerea, Aspicilia* sp., *S. testudinea* and *R. geographicum)* of lichens’ presence, year of exposure, and the interaction of presence and year of exposure.

|  | Df | F value | P value |
| --- | --- | --- | --- |
| Lichen Presence | 1 | 14.5 | < 0.001 |
| Year | 4 | 5.8 | 0.002 |
| Lichen Presence x Year | 4 | 2.6 | 0.061 |

## Discussion

The overall aim of this study was to identify whether different lichen species and growth forms affect rock surface weathering. Our findings confirm that all three growth forms were associated with reduced rock hardness and, given that declines in rock hardness are associated with increased surface weathering in Antarctica (Guglielmin et al. 2012, Kanamaru et al. 2018), provide support for our hypothesis that crustose lichens have a greater impact on rock weathering than foliose and fruticose lichens (Fig. 1). Furthermore, our results demonstrate that lichen impacts on rock weathering increase with surface exposure time and lichen thallus size, consistent with the hypothesis that older and larger lichens have a greater effect on rock hardness (Fig. 5). These findings provide new insights into the contributions of different lichen growth forms to weathering of rock surfaces, expanding on previous work focused primarily on crustose lichens. Growth form is an important trait in lichens with functional consequences for their local ecosystems (Asplund and Wardle 2017). The impacts of crustose lichens in this context have been studied along the western Antarctic Peninsula (McCarroll and Viles 1995; Matthews and Owen 2008; Guglielmin et al. 2012), but fruticose and foliose lichens have received little research attention (Chen et al. 2000), despite also developing on rock substrata and having extensive distributions in this region (Holdgate 1977; Peat et al. 2007).

### Variation in rock hardness among lichen growth forms

Our data confirm that, generally, the crustose lichens studied had the strongest influence on rock substrate hardness, as reported in previous studies (Wilson and Jones 1983). However, our data also indicate that some crustose species may have much weaker effect than hypothesized or even none at all, as illustrated by *R. geographicum*. Following a chronosequence approach as described by McCarroll and Viles (1995) and Matthews and Owen (2008), we assessed the influence of four lichen species on rock hardness over time in the Morteratsch Glacier foreland in the Swiss Alps. Amongst these four species, there were large differences in the rates of weathering documented, even though the overall rock weathering effect was positive in the absence of any confounding effects of lichens colonising pre-weathered rock surfaces (see below). It is clear from our data that knowledge of the identity of individual lichen species is required to provide an accurate assessment of their influence on rock weathering.

Although crustose lichens generally had the strongest effects on surface weathering, our data also confirm that foliose and fruticose lichens had measurable impacts on rock hardness. The larger impact of crustose lichens most likely derives from greater surface contact area, allowing for higher hyphal density penetrating the rock surface (Chen et al. 2000). Hyphal growth can fracture rock surfaces by the repeated expansion and shrinking of hyphae (Adamo and Violante 2000). Once established, these hyphae allow water entry into micro fissures in the rock which, in polar regions, can result in ice expansion during repeated freeze-thaw cycles, contributing further to fracturing the rock surface. This process is also likely to take place around the anchoring points of foliose and fruticose lichens, but focused in a much smaller surface area (Fig. 6). Foliose lichens have been associated with surface weathering, but mainly in softer rock types (Adamo and Violante 2000). Foliose lichens have also been suggested to have hyphae that penetrate deeper into the rock surface, resulting in greater weathering than observed for fruticose lichens (De Los Rios et al. 2005, Guglielmin et al. 2012), and this is reflected in our data. However, there were clear species-specific differences in rock hardness effects, cautioning that the categorical growth form variation may also obscure relevant species-specific traits. The miniature fruticose lichen *P. minuscula,* for instance, did not affect rock hardness detectably, and the crustose *R. geographicum* had a much weaker influence than other crustose species. Further research should elucidate the traits that determine such interspecific variation in weathering capacity. Overall, while our data support crustose lichens having a stronger influence on rock hardness than foliose and fruticose species, given the extensive cover of these groups in fellfield across habitats in many parts of the sub- and maritime Antarctic regions (Holdgate 1977, Smith 1995, Øvstedal and Smith 2001), the accumulated effect of many million individual fruticose lichens may still contribute substantially to large-scale weathering processes.

**Fig. 6.**
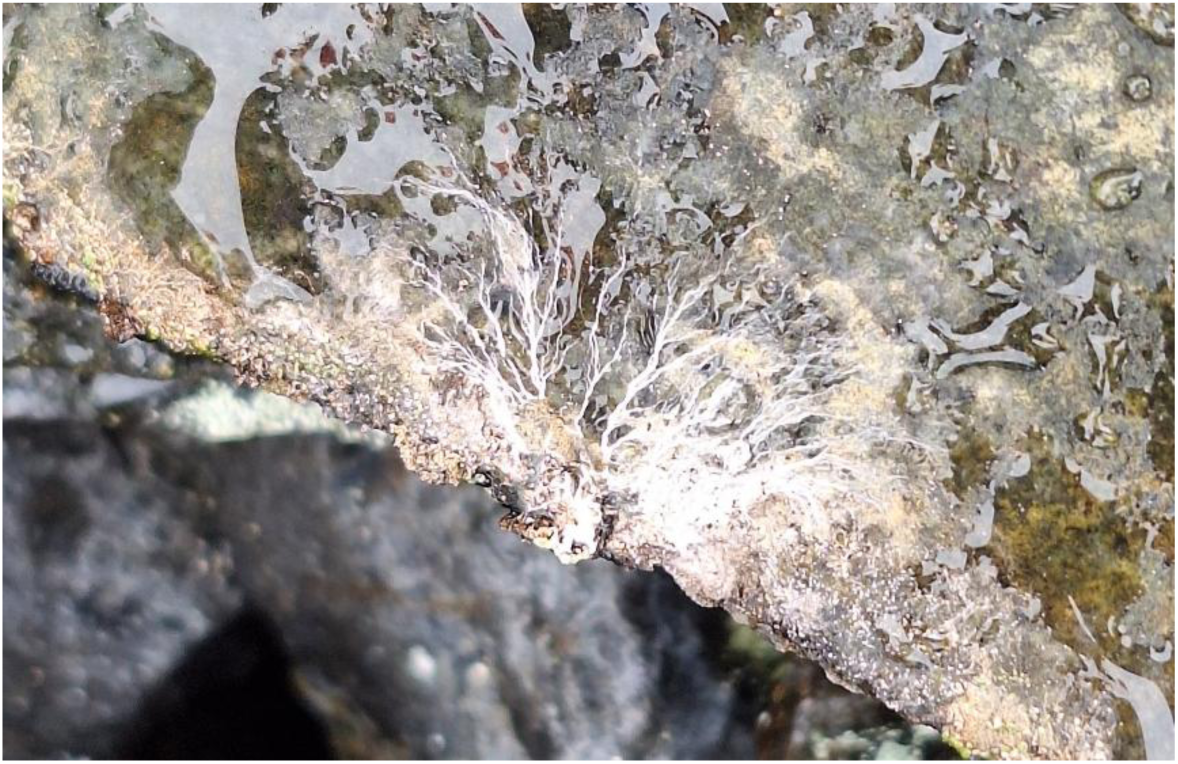
An example of a fruticose lichen’s anchoring point. Photo taken by Peter Convey near Rothera Station on Leonie Island.

### Solving the chicken-and-egg problem of causality in rock weathering by lichens

Our data also provided strong support for our second hypothesis, that lichen impacts on rock hardness are an active, biologically driven, process that increases with rock surface age, leading to a significantly stronger impact on rock surfaces that have been exposed for longer than on those more recently exposed. While bare rock naturally weathers over time due to various factors (Wilson and Jones 1983), the higher rates of weathering on lichen-encrusted surfaces observed in the Swiss Alps are clearly consistent with this hypothesis. Lichens actively contribute to mineral dissolution and breakdown, likely through a combination of mechanical and chemical weathering by hyphal penetration and acid production (Chen et al. 2000). The data obtained in the Swiss Alps confirm that weathering was increased when the lichen species, *A. cinerea, S. testudinea* and *Aspicilia* sp., were present. This observation is consistent with other studies that have modelled crustose lichen growth (Armstrong 2011), showing that crustose lichens experience distinct growth phases, with an initial lag, followed by accelerated growth and, finally, a slow phase (Armstrong 1983; Armstrong and Bradwell 2010). During periods of growth, lichen metabolic activity can be very high (Chen et al. 2000). This suggests that older lichen thalli may contribute less actively to weathering due to reduced metabolic rates, again consistent with our data.

Our third hypothesis was that lichen size (as a proxy for lichen age as well as an indication of time since deglaciation) would be correlated with a greater reduction in rock hardness over time. In the Swiss Alps, all four lichen species examined were generally characterised by softer substrate beneath larger lichen specimens. This is consistent with older, more established, lichens having already increased the rate of rock surface weathering, and also aligns with the outcome of our second hypothesis, as discussed above, which suggests that the growth phase of individual lichen thalli and the resulting metabolic activity as the thallus grows will influence the rate of rock surface weathering. The margins of a growing lichen thallus will contract and expand slightly with the loss or gain of moisture (Benedict 2008), which will provide chronic pressure on the underlying substratum. Larger lichen thalli can retain greater amounts of moisture, which could cause a positive feedback where a micro-environment is formed that enhances rock degradation. The relationships between thallus size and rock surface hardness were statistically significant for *R. geographicum, Aspicilia* sp. and *S. testudinea*, but not for *A. cinerea*, again highlighting that inferences based on one species cannot automatically be extended to all lichen species. Variation between microhabitats is a critical influence on lichen growth rates (Armstrong 2015) and is likely to influence rock weathering effects.

*Rhizocarpon geographicum* may be an outlier in both the Antarctic and Swiss datasets because the rock it grew on was inherently softer than that colonised by the other species studied here. This is consistent with observations that this species has a faster growth on softer rock types (Armstrong 2011). It may require softer, already weathered, rock surfaces for establishment, contrasting with species such as *Buellia frigida* and *Aspicilia* sp., which were often present on surfaces that had very recently been deglaciated. All three of these genera are typically found close to glacier forefronts and in moraines in the polar regions (Wietrzyk et al. 2017), where they are amongst the first colonizers of recently exposed substrata. As lichens also retain moisture more easily than bare rock (Asplund and Wardle 2017), it is possible that the increased moisture availability makes certain substrata more preferable for *R. geographicum* colonization. Other studies have reported that *R. geographicum* tends to have higher growth rates where there is increased moisture availability (Bradwell 2001, Matthews 2005).

### Outlook and conclusions

Various other aspects of potential interactions between lichens and their rock substrate should receive attention in future research. For instance, crustose lichen species commonly abut to each other and even fuse over time, or with species of other lichen growth forms, forming often complex mosaics (Armstrong and Bradwell 2010) that may behave differently to individual lichen species. These were excluded in the current study to avoid introducing further confounding factors. Future research should also consider the sub-growth forms of lichen morphology, such as areolate, leprose and placodoid lichens, all of which are variations of crustose lichens (Untari 2024). The sections of an areolate thallus and the undifferentiated, powdery, thallus of a leprose lichen clearly differ morphologically, such that their effects on rock weathering may differ considerably. Fractured or crumbling rock surfaces were also not considered in the current study, but we recognise that this crumbling could itself be the result of lichen-induced weathering that has weakened the rock surfaces to such an extent that it fractures easily when struck.

Overall, our study supports the hypothesis that rock hardness beneath lichens differs with both the overgrowing lichen species and their growth form, and highlights that all major growth forms should be considered in studies of lichen influence on rock weathering. Interactions between these traits are complex and differ between individual lichen species. In-depth future studies to reveal the traits and mechanisms underpinning such interspecific variation will be important for better understanding rock weathering processes in cold biomes.

## Declarations

The authors have no competing interests to declare that are relevant to the content of this article. This research was part of the Antarctic Biota Count project (ALWPP.2019.006), which was funded by the Netherlands Organisation for Scientific Research (NWO). P. Convey is supported by NERC core funding to the British Antarctic Survey’s ‘Biodiversity, Evolution and Adaptation’ Team.

## Acknowledgements

This research was part of the Antarctic Biota Count project (ALWPP.2019.006), which was funded by the Netherlands Organisation for Scientific Research (NWO). P. Convey is supported by NERC core funding to the British Antarctic Survey’s ‘Biodiversity, Evolution and Adaptation’ Team. The authors wish to thank staff at the BAS Rothera Station for practical and logistical support.

## Author Contributions

E. N. Ciric wrote this manuscript and participated in field work and sample collection. I.K. de Jonge assisted in sample collection and statistics. R. Liu assisted in statistics and editing. J. H. C. Cornelissen supervised this project and assisted in editing the manuscript. P. Convey supervised this project and assisted in editing the manuscript. S. Bokhorst participated in field work, sample collection, supervision, and did statistics and editing on the manuscript.

## Supplementary Material

**SI Table 1.**
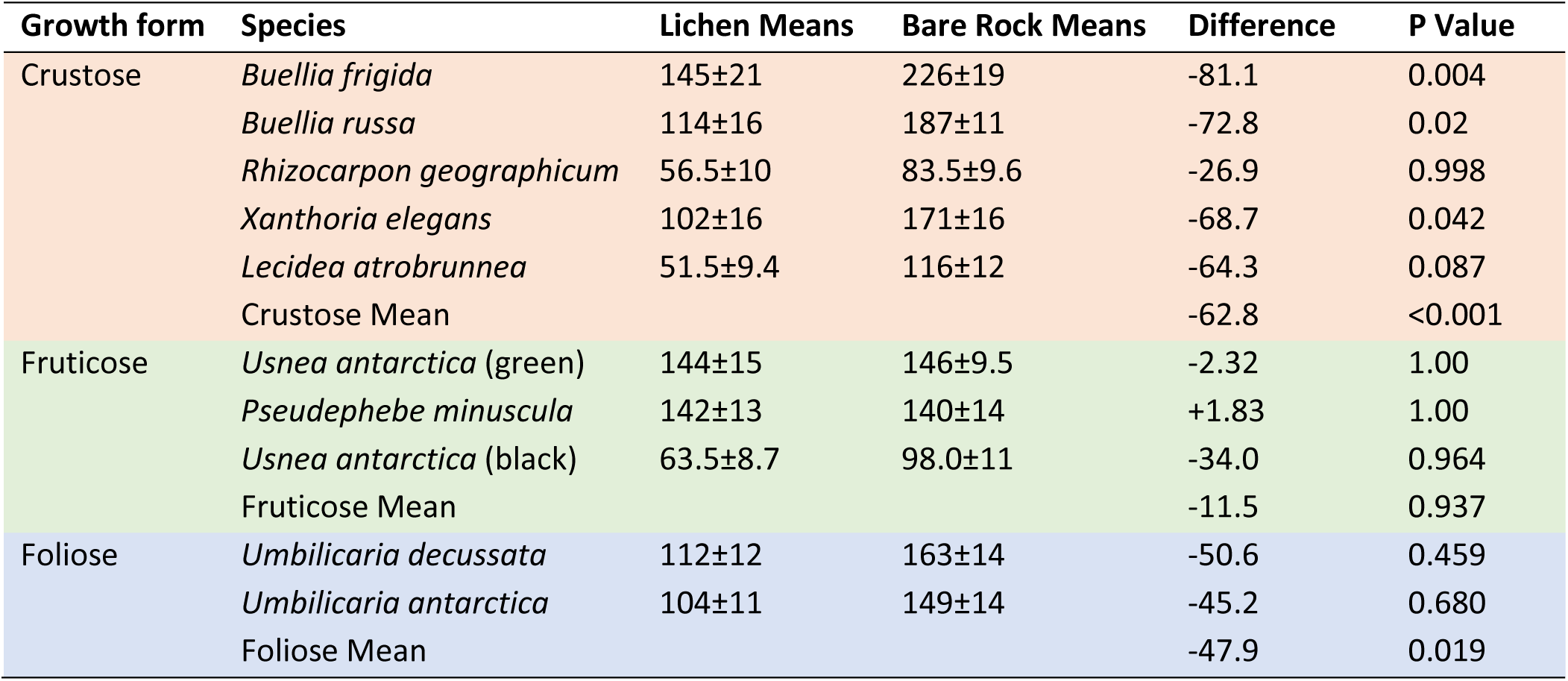
Lichen species effects on rock hardness in Antarctica, showing the mean rock hardness of 10 Antarctic lichen species and neighbouring bare rock without lichen presence. Values are means of n = 25 with SE in parentheses. P-values were obtained from Tukey HSD tests to identify the effect of lichen presence on rock hardness.

| Growth form | Species | Lichen Means | Bare Rock Means | Difference | P Value |
| --- | --- | --- | --- | --- | --- |
| Crustose | *Buellia frigida* | 145±21 | 226±19 | -81.1 | 0.004 |
|  | *Buellia russa* | 114±16 | 187±11 | -72.8 | 0.02 |
|  | *Rhizocarpon geographicum* | 56.5±10 | 83.5±9.6 | -26.9 | 0.998 |
|  | *Xanthoria elegans* | 102±16 | 171±16 | -68.7 | 0.042 |
|  | *Lecidea atrobrunnea* | 51.5±9.4 | 116±12 | -64.3 | 0.087 |
|  | Crustose Mean |  |  | -62.8 | <0.001 |
| Fruticose | *Usnea antarctica* (green) | 144±15 | 146±9.5 | -2.32 | 1.00 |
|  | *Pseudephebe minuscula* | 142±13 | 140±14 | +1.83 | 1.00 |
|  | *Usnea antarctica* (black) | 63.5±8.7 | 98.0±11 | -34.0 | 0.964 |
|  | Fruticose Mean |  |  | -11.5 | 0.937 |
| Foliose | *Umbilicaria decussata* | 112±12 | 163±14 | -50.6 | 0.459 |
|  | *Umbilicaria antarctica* | 104±11 | 149±14 | -45.2 | 0.680 |
|  | Foliose Mean |  |  | -47.9 | 0.019 |

**SI Table 2.** Results of T-tests to identify which lichen species have significant effects on rock weathering based on LRR data.

| Species | P Value | Mean LRR |
| --- | --- | --- |
| *Buellia frigida* | 0.001 | -0.633 |
| *Buellia russa* | < 0.001 | -0.071 |
| *Xanthoria elegans* | < 0.001 | -0.665 |
| *Lecidea atrobrunnea* | < 0.001 | -1.090 |
| *Rhizocarpon geographicum* | 0.004 | -0.501 |
| *Usnea antarctica* (green) | 0.20 | -0.113 |
| *Pseudephebe minuscula* | 0.59 | 0.032 |
| *Usnea antarctica* (black) | < 0.001 | -0.549 |
| *Umbilicaria decussata* | 0.009 | -0.431 |
| *Umbilicaria antarctica* | 0.006 | -0.409 |

**SI Table 3.** Results of ANCOVA with a linear model function with log response ratio as the dependent variable, and species, years of exposure and thallus size as the predictors.

| Variable | Estimate | Std. Error | t value | p value |
| --- | --- | --- | --- | --- |
| *Aspicilia* sp. | -0.13 | 0.07 | -1.84 | 0.07 |
| *S. testudinea* | 0.14 | 0.05 | 2.65 | 0.008 |
| *A. cinerea* | -0.03 | 0.05 | -0.49 | 0.63 |
| *R. geographicum* | -0.24 | 0.06 | -4.35 | < 0.001 |
| Year: 1896 | 0.23 | 0.06 | 3.66 | < 0.001 |
| Year: 1917 | 0.06 | 0.07 | 0.98 | 0.33 |
| Year: 1930 | -0.02 | 0.06 | 0.32 | 0.75 |
| Year: 1955 | 0.05 | 0.07 | 0.69 | 0.49 |
| Year: 1971 | 0.18 | 0.09 | 1.99 | 0.05 |
| Year: 1998 | 0.19 | 0.12 | 1.61 | 0.11 |
| Size | -0.01 | 0.001 | -7.02 | < 0.001 |

**SI Fig.1.**
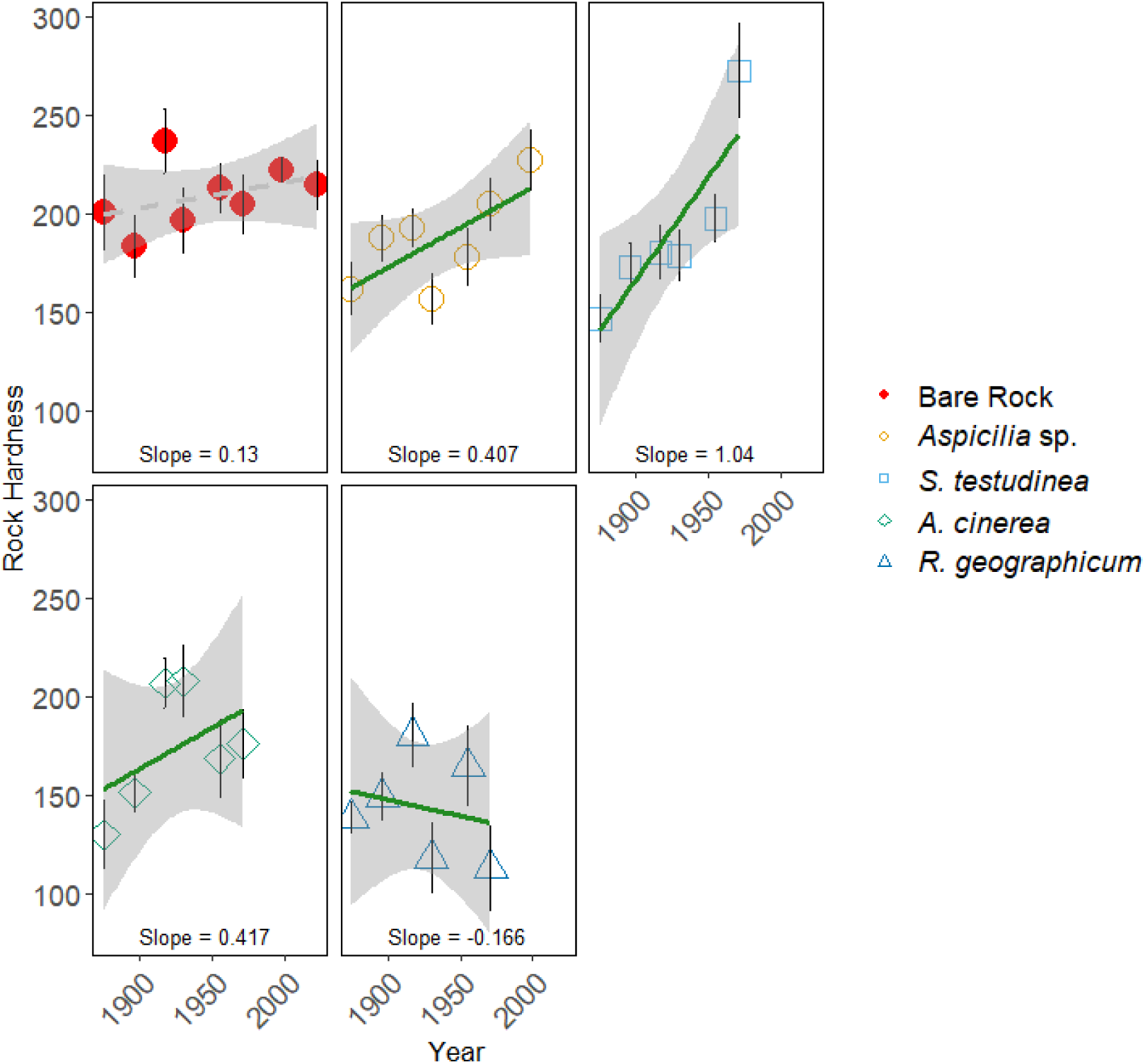
The hardness of bare rock and each individual lichen species studied in Switzerland along a dated chronosequence of glacial retreat. Trend lines indicate the change in rock hardness over time for bare rock (dotted line) and beneath lichens (solid line). Symbols are the mean of 10 replicate sampling sites at each chronosequence stage (with SE as error bars). Ninety-five percent confidence intervals are represented by grey shading.

